# Differential effects of two catalytic mutations on full-length PRDM9 and its isolated PR/SET domain reveal a case of pseudo-modularity

**DOI:** 10.1101/2021.08.09.455705

**Authors:** Natalie R. Powers, Timothy Billings, Kenneth Paigen, Petko M. Petkov

**Affiliations:** The Jackson Laboratory, Bar Harbor, Maine, 04609, USA

**Keywords:** PRDM9, PR/SET domain, point mutations, enzyme activity, infertility phenotype

## Abstract

PRDM9 is a DNA-binding histone methyltransferase that designates and activates recombination hotspots in mammals by locally trimethylating lysines 4 and 36 of histone H3. In mice, we recently reported two independently produced point mutations at the same residue, glu360pro (*Prdm9*^EP^) and glu360lys (*Prdm9*^EK^), which severely reduce its H3K4 and H3K36 methyltransferase activities *in vivo. Prdm9*^EP^ is slightly less hypomorphic than *Prdm9*^EK^, but both mutations reduce both the number and amplitude of PRDM9-dependent H3K4me3 and H3K36me3 peaks in spermatocytes. While both mutations cause infertility with complete meiotic arrest in males, *Prdm9*^EP^, but not *Prdm9*^EK^, is compatible with some female fertility. When we tested the effects of these mutations *in vitro*, both *Prdm9*^EP^ and *Prdm9*^EK^ abolished H3K4 and H3K36 methyltransferase activity in full-length PRDM9. However, in the isolated PRDM9 PR/SET domain, these mutations selectively compromised H3K36 methyltransferase activity, while leaving H3K4 methyltransferase activity intact. The difference in these effects on the PR/SET domain versus the full-length protein show that PRDM9 is not an intrinsically modular enzyme; its catalytic domain is influenced by its tertiary structure and possibly by its interactions with DNA and other proteins *in vivo*. These two informative mutations illuminate the enzymatic chemistry of PRDM9, and potentially of PR/SET domains in general, reveal the minimal threshold of PRDM9-dependent catalytic activity for female fertility, and potentially have some practical utility for genetic mapping and genomics.

## Introduction

Genetic recombination, the process by which genetic information is exchanged between homologous chromosomes, is both an important source of genetic diversity and a necessary step for successful meiosis in most organisms. In many taxa, recombination events are restricted to specific sites in the genome known as recombination hotspots. Many mammalian species, including mice and humans, utilize the specialized histone methyltransferase PRDM9 to determine the locations of hotspots and to activate them for the recombination process (Paigen and Petkov 2018). In the PRDM9-mediated system, hotspots are restricted to the cognate binding sites of PRDM9 and subsequently marked by locally trimethylating lysines 4 and 36 of histone H3 (Hayashi *et al*. 2005; Baudat *et al*. 2010; Berg *et al*. 2010; Parvanov *et al*. 2010; Baker *et al*. 2014; Powers *et al*. 2016). This double histone mark creates a unique epigenetic signature that differentiates hotspots from H3K4me3-enriched promoters and regulatory elements, and from the H3K36me3-enriched bodies of actively transcribed genes.

PRDM9 was previously thought to be necessary for successful meiosis in mammals, largely because C57BL/6J (B6) *Prdm9* knock-out mice are infertile with meiotic arrest (Hayashi *et al*. 2005), and also because *PRDM9* has been associated with azoospermia in humans (Miyamoto *et al*. 2008; Irie *et al*. 2009). The catalytic activity of PRDM9 was likewise considered necessary for fertility, as B6 mice homozygous for a methyltransferase-dead *Prdm9* allele are also infertile (Diagouraga *et al*. 2018). However, exceptions to this rule have accumulated since the discovery of *Prdm9*. PWD/Ph male mice produce small amounts of mature sperm in the absence of *Prdm9* (Mihola *et al*. 2019). Canines manage to carry out meiosis without impediment, despite pseudogenization of *Prdm9* in that lineage (Munoz-Fuentes *et al*. 2011; Auton *et al*. 2013). And notably, there is at least one case of a fertile *PRDM9*-null human female (Narasimhan *et al*. 2016).

The structure of PRDM9 appears to be modular: it has three well-defined and well-separated functional domains—KRAB, PR/SET, and a tandem zinc finger (ZnF) array. The KRAB domain is involved in protein-protein interactions (Imai *et al*. 2017; Parvanov *et al*. 2017; Diagouraga *et al*. 2018). The PR/SET domain uniquely trimethylates histone H3 at both lysines 4 and 36 (Hayashi *et al*. 2005; Baker *et al*. 2014; Powers *et al*. 2016). The ZnF domain binds DNA in a sequence-specific manner (Brick *et al*. 2012; Billings *et al*. 2013; Walker *et al*. 2015; Smagulova *et al*. 2016). Consistent with its apparently modular structure, the PRDM9 catalytic PR/SET domain and its DNA-binding ZnF domain retain their functions when isolated from the rest of the protein *in vitro*, suggesting that these domains are autonomous (Billings *et al*. 2013; Wu *et al*. 2013; Eram *et al*. 2014; Koh-Stenta *et al*. 2014; Walker *et al*. 2015; Powers *et al*. 2016). Inactivating mutations in either domain cause sterility in mutant mice, emphasizing the importance of each domain for the overall function of PRDM9 (Parvanov *et al*. 2017; Diagouraga *et al*. 2018).

In mice, we recently reported sexual dimorphism in the requirement for PRDM9 for successful meiosis (Powers *et al*. 2020). While both male and female B6 and C3H/HeJ (C3H) mice are infertile in the absence of *Prdm9*, female *Prdm9*-null CAST/EiJ (CAST) mice are fully fertile and produce healthy offspring, although male *Prdm9*-null CAST mice are infertile. We also reported two catalytic mutations—*Prdm9*^EP^ and *Prdm9*^EK^—that straddle the threshold for female fertility in B6 mice (Bhattacharyya *et al*. 2019; Powers *et al*. 2020). These mutations substitute glutamate 360, a residue situated within the PR/SET domain substrate-binding groove, with proline and lysine, respectively. *In vivo*, both mutations result in a catalytically hypomorphic PRDM9 protein that is aptly described as ‘methyltransferase-comatose’; both H3K4 and H3K36 methyltransferase activities are severely reduced, but not completely abolished. *Prdm9*^EK^ encodes a slightly less active protein than *Prdm9*^EP^, and this results in a sex-specific fertility phenotype. Males homozygous for either mutation exhibit a phenocopy of the *Prdm9*-null and *Prdm9-*methyltransferase-dead conditions: infertility with complete meiotic arrest in the pachytene stage of prophase I. Homozygous female *Prdm9*^*EK*^ mice are also infertile (Bhattacharyya *et al*. 2019). In contrast, homozygous female *Prdm9*^EP^ mice are able to produce variable numbers of grossly healthy, normal offspring via natural matings (Powers *et al*. 2020). These observations show that oogenesis is robust to less PRDM9-dependent catalytic activity than spermatogenesis, even on the B6 genetic background.

In this paper, we characterize and contrast *Prdm9*^EP^ and *Prdm9*^EK^, and present experimental data demonstrating the disparate effects of these mutations on full-length PRDM9 versus its isolated PR/SET domain. We show that both mutations severely reduce or abolish the catalytic activity of full-length PRDM9 *in vivo* and *in vitro*. However, the two mutations show separation-of-function in the context of the PR/SET domain alone *in vitro*. Recombinant *Prdm9*^EP^ and *Prdm9*^EK^ PR/SET domains show largely uncompromised H3K4 trimethylation activity, while H3K36 trimethylation activity is reduced by the *Prdm9*^EK^ mutation and abolished by the *Prdm9*^EP^ mutation. This observation shows that PRDM9 is not truly modular, as *Prdm9*^EP^ and *Prdm9*^EK^ have different effects on the full-length protein vs. the isolated PR/SET domain. We discuss the implications of these mutations for different aspects of PRDM9 function— enzyme chemistry, the differential importance for female vs. male meiosis, and its evolution.

## Materials and Methods

### Ethics statement

The animal care rules used by The Jackson Laboratory are compatible with the regulations and standards of the U.S. Department of Agriculture and the National Institutes of Health. The protocols used in this study were approved by the Animal Care and Use Committee of The Jackson Laboratory (summary # 04008). Euthanasia was carried out by cervical dislocation.

### Mouse Strains

Mice used in this study were acquired from The Jackson Laboratory. The following strains were used: C57BL/6J (stock number 000664), C57BL/6J-*Prdm9*^<em2Kpgn>^/Kpgn (*Prdm9*^EP^), and C57BL/6J-*Prdm9*^<em2Kpgn>^/Kpgn (*Prdm9*^EK^). The latter two were generated by introducing a point mutation (Glu360Pro) via CRISPR-Cas9 gene editing, onto the C57BL/6J genetic background, as previously described (Bhattacharyya *et al*. 2019; Powers *et al*. 2020). Gene editing was performed by the Genetic Engineering Technology core at The Jackson Laboratory.

### Fertility Tests

Each *Prdm9*^EK/EK^ female (5 weeks old) was housed with fertile, sexually mature (>8 weeks old) wild-type B6 male for a period of at least three and up to seven months. A female was considered fertile if she gave live birth at least once. Litter size was determined by counting pups on the day of birth.

### PRDM9 PR/SET Domain Synthesis and Cloning

The genetic sequence encoding amino acids 187-372 of the mouse reference sequence of PRDM9 was synthesized and cloned into the pBAD-HisB expression vector, using the GenScript commercial service. In the case of *Prdm9*^EP^ and *Prdm9*^EK^, the codon for glutamate 360 was altered to encode proline and lysine, respectively. Sanger sequencing confirmed that the sequences were correct, and that there were no off-target mutations.

### Site-Directed Mutagenesis and Cloning of Full-length PRDM9

To induce the *Prdm9*^EP^ and *Prdm9*^EK^ mutations in full-length PRDM9, we used site-directed mutagenesis with a wild-type cDNA version of human *PRDM9* (allele C) that we had previously cloned into pBAD-HisB. We performed site-directed mutagenesis using the QuikChange II kit (Agilent), as per the manufacturer’s instructions. Sanger sequencing confirmed that the sequences were correct, and that there were no off-target mutations.

### Protein Expression

To facilitate expression and purification, the sequences encoding both the PR/SET domains and full-length PRDM9 were transferred into the pMAL-c6T (New England Biolabs #N0378) vector, which contains a maltose-binding (MBP) tag. The constructs were transformed into NEB stable cells (NEB C3040I). Plasmid DNA was obtained from individual colonies and sequenced to find the correct clone. Plasmids were then transformed into Rosetta II cells (Sigma -71402-4) and grown on agar plates with 100mg/ml ampicillin overnight at 37°C.

Single colonies of each clone were used for overnight expression culture in 100 ml in LB with 100mg/ml ampicillin at 30°C. No inducer was added because expression is leaky and allows production of enough protein for further analysis.

### Purification and Histone Methyltransferase (HMT) Assays

Cell lysis, protein isolation, and HMT assays were done as previously described (Powers *et al*. 2016). Briefly, cells were lysed in 1xCBB (cell breakage buffer), consisting of 500 mM Tris-HCl pH 7.5, 0.15M sucrose, 0.1% NP-40, 100 mM KCl and Roche EDTA-free complete protein inhibitors (cat. #04693132001) and sonicated for 1-2 minutes on ice. The supernatant was cleared by centrifugation at 12,000xg for 20 minutes. The expressed proteins were partially purified on Q sepharose (Sigma q1126) and the fractions containing the protein of interest were collected. Western blotting was done using an iBlot dry transfer system, as per manufacturer’s instructions. Primary antibodies are identified and used in concentrations as follows: α-H3K4me3: 1:5000, EMD Millipore cat#07–473; α-H3K36me3: 1:5000, Active Motif cat#61101; α-Histone H3: 1:5000, Active Motif cat#61277. The secondary antibody used with all three primary antibodies was HRP-conjugated goat anti-rabbit (Bio-Rad cat#1662408EDU). Blots were imaged on a SynGene gel imaging system after a 1min exposure to SuperSignal Femto West Maximum Sensitivity Substrate (ThermoFisher), and band density was quantified using GeneTools software. Three replicate HMT assays were done for each PR/SET domain and full-length PRDM9 protein; their means and standard deviations are plotted in Figures 1 and 2.

**Figure 1:**
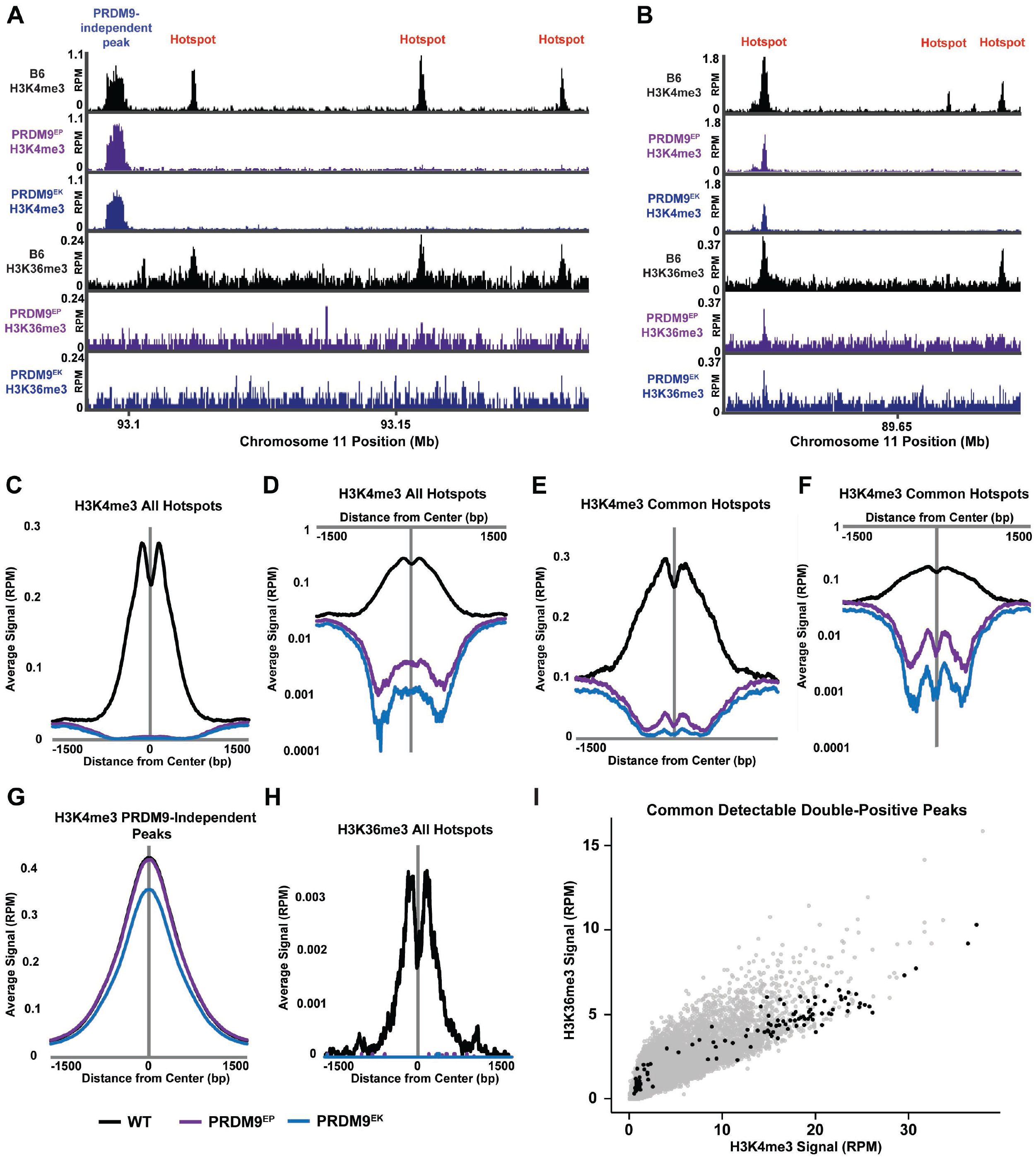
Histone methyltransferase activities of wild-type, PRDM9^EP^, and PRDM9^EK^ PR/SET domains. (**A-C**) Representative western blots showing the results of histone methyltransferase (HMT) assays. Blots were probed with α-H3K4me3 and α-H3K36me3 antibodies to show the levels of these modified histones at the denoted time points (minutes) as the assay proceeded. Negative control lanes (-) represent reactions run without s-adenosyl methionine (SAM), while positive control lanes (+) represent purified H3K4me3 or H3K36me3, as appropriate. H3K4me3 blots were stripped and re-probed with α-histone H3 antibody to confirm the presence of histone at all time points. (**D-E)** Time course plots summarizing HMT activities for H3K4me3 (**D**) and H3K36me3 (**E**), quantified by band densitometry. Data points represent the mean of three replicates; error bars represent the standard deviations.

**Figure 2:**
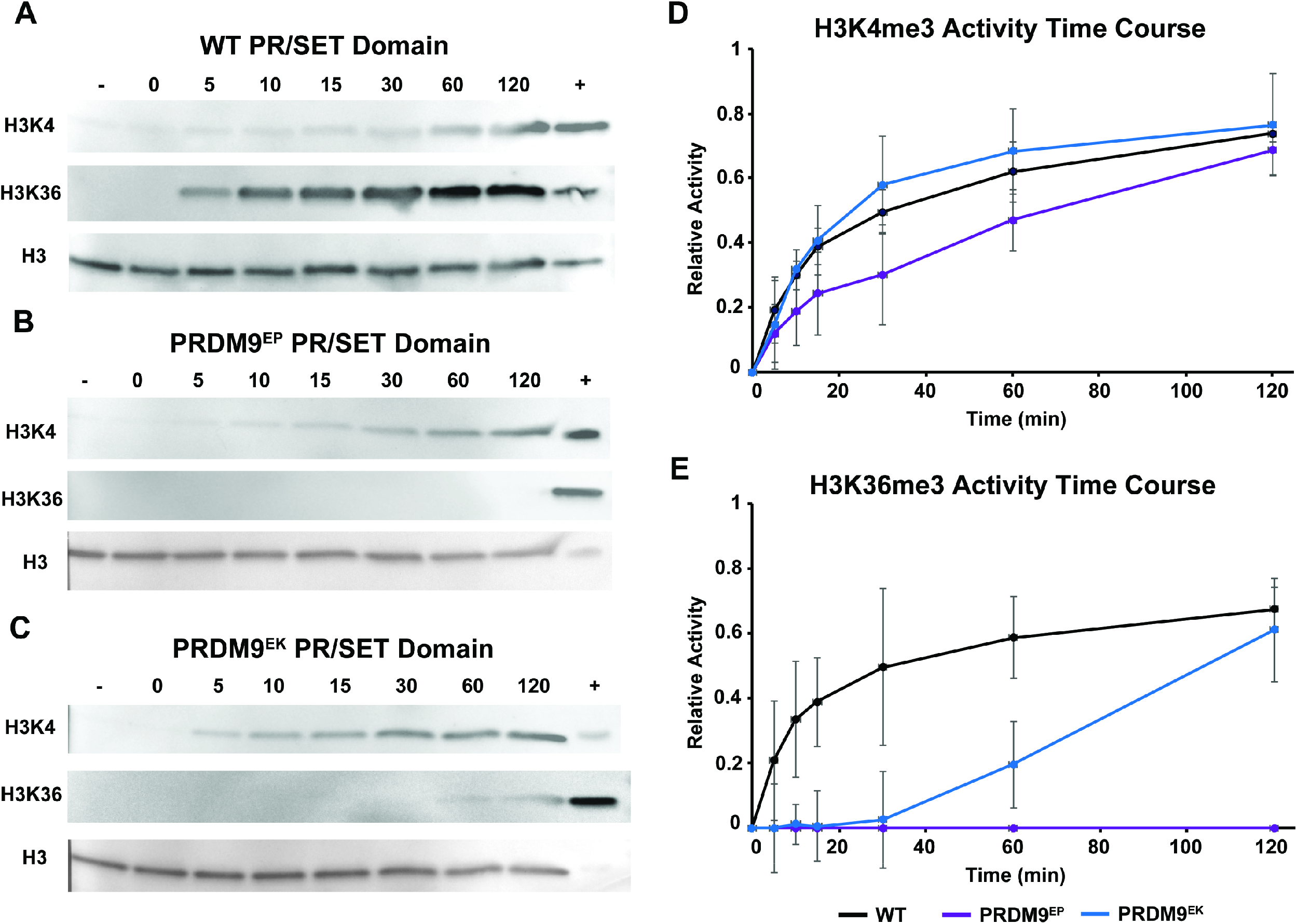
Histone methyltransferase activities of full-length wild-type, PRDM9^EP^, and PRDM9^EK^. (**A-C**) Western blots showing the results of histone methyltransferase (HMT) assays with full-length proteins. Blots were probed with α-H3K4me3 and α-H3K36me3 antibodies to show the levels of these modified histones at the denoted time points (minutes) as the assay proceeded. Negative control lanes (-) represent reactions run without s-adenosyl methionine (SAM), while positive control lanes (+) represent purified H3K4me3 or H3K36me3, as appropriate. H3K4me3 blots were stripped and re-probed with α-histone H3 antibody to confirm the presence of histone at all time points. (**D-E)** Time course plots summarizing HMT activities for H3K4me3 (**D**) and H3K36me3 (**E**), quantified by band densitometry. Data points represent the mean of three replicates; error bars represent the standard deviations.

### H3K4me3 ChIP-seq

ChIP-seq for H3K4me3 was performed using spermatocytes isolated from 14-day-old males, as described previously (Baker *et al*. 2014). For B6 and *Prdm9*^EP/EP^ spermatocytes, ChIP-seq data our lab previously reported (Baker *et al*. 2014; Powers *et al*. 2020) was used. In both cases, two biological replicates were merged for analysis, with a total of 39,973,027 aligned reads for B6 and 45,931,625 aligned reads for *Prdm9*^EP/EP^ spermatocytes, as in Powers et al. (2020). For *Prdm9*^EK/EK^ spermatocytes, one replicate was previously reported (Bhattacharyya *et al*. 2019), and an additional biological replicate was performed for this study, for a total of two replicates per genotype. This additional replicate was sequenced on the Illumina NextSeq 500 platform, with 75-bp reads, and trimmed for quality using trimmomatic. Sequence data were aligned to the mouse mm10 genome using BWA (Burrows-Wheeler aligner) v0.7.9a, and duplicate reads and reads which failed to align to unique positions in the genome were discarded. For the additional *Prdm9*^EK/EK^ replicate, this resulted in 56,096,669 aligned reads. The raw fastq file of this replicate was combined with that of the previous replicate for analysis, and the merged file was aligned to mm10 via the procedure above for a total of 72,730,785 aligned reads.

H3K4me3 peaks were called using MACS v1.4, default parameters, with a *P* value cutoff of 0.01, using treatment and control samples. The same procedure and samples that we used in our previous study (Powers *et al*. 2020) were used, with the addition of the merged *Prdm9*^EK/EK^ sample, as well as the set of PRDM9-dependent H3K4me3 peaks described in that study. To determine overlap between known PRDM9-dependent H3K4me3 peaks and ChIP-seq peak sets, we used BedTools (v2.27.0) intersect with default parameters, except for a requirement for 20% overlap (*f* = 0.20).

### H3K36me3 ChIP-seq

ChIP-seq for H3K36me3 was performed using spermatocytes isolated from 14-day-old males, as described previously (Powers *et al*. 2016). One biological replicate per genotype was performed. DNA samples were sequenced on an Illumina HiSeq 2500, with 100bp reads, and trimmed for quality using trimmomatic. Sequence data were aligned to the mouse mm10 genome using BWA (Burrows-Wheeler aligner) v0.7.9a, and duplicate reads and reads which failed to align to unique positions in the genome were discarded. This resulted in a total of 42,288,943 aligned reads for *Prdm9*^EP/EP^ spermatocytes, and 40,263,687 aligned reads for *Prdm9*^EK/EK^ spermatocytes. For the B6 control, we used two previously reported biological replicates (Powers *et al*. 2016), merging the raw fastq files and aligning to the mm10 genome for analysis. This resulted in 95,375,802 aligned reads.

To assess detectable H3K36me3 peaks, we also used MACS v1.4, default parameters, with a *P* value cutoff of 0.01, but in this case with treatment samples only so that enrichment for each peak was determined relative to the surrounding background, rather than relative to a control sample. We chose this method because H3K36me3 is broadly enriched in transcriptionally active genes, and this broad enrichment can confound peak calling relative to a control sample. To determine overlap between known PRDM9-dependent H3K4me3 peaks and ChIP-seq peak sets, we used BedTools (v2.27.0) intersect with default parameters, except for a requirement for 20% overlap (*f* = 0.20).

### Aggregation Plots

Aggregation plots were done using the Aggregation and Correlation Toolbox (ACT, http://act.gersteinlab.org/), with the following parameters: nbins = 500, mbins = 0, radius = 1500.

## Results

### *Prdm9*^EP^ and *Prdm9*^EK^ mutations compromise the H3K36 trimethylation activity of the isolated PRDM9 PR/SET domain *in vitro*

In wild type mice, double-stranded breaks (DSBs) that initiate recombination occur at PRDM9-activated sites and their numbers are tightly controlled. Only about 5% of PRDM9-bound sites undergo DSB formation (Kauppi *et al*. 2013). In *Prdm9*-null B6 mice, DSBs are made in normal numbers, but at ectopic sites where H3K4me3 is enriched, such as promoters. These ectopic DSBs are not repaired efficiently and may be the principal cause of the meiotic arrest followed by apoptosis due to loss of *Prdm9* (Brick *et al*. 2012). Presumably, this specific meiotic effect could be ascribed to the unique activity of the PR/SET domain of PRDM9, which combines two methyltransferase activities—trimethylation of histone H3 at both lysine 4 and lysine 36 (Powers *et al*. 2016). To further study the nature of this activity and its effect on DSB initiation and repair, we created two mutants, *Prdm9*^EP^ and *Prdm9*^EK^.

We studied the effects of these mutations on PR/SET domain methyltransferase activities by performing histone methyltransferase (HMT) time-course assays using wild-type, *Prdm9*^EP^ and *Prdm9*^EK^ PR/SET domains, with three replicates of each time course. Figure 1A-C shows representative blots, and Figure 1D-E shows time course plots with the average and standard deviation of the three replicates. The H3K4 trimethylation curves of *Prdm9*^EP^ and *Prdm9*^EK^ PR/SET domains were indistinguishable from the one of wild-type PR/SET domain (Fig. 1D). In contrast, the H3K36 methyltransferase activities of the mutant PR/SET domains were substantially different from the wild-type domain (Fig. 1E). The initial rate of the H3K36 trimethylation activity of *Prdm9*^EK^ PR/SET domain was substantially reduced, though the average did reach wild-type levels of H3K36 trimethylation at the 120-minute endpoint. For the *Prdm9*^EP^ PR/SET domain, H3K36 trimethylation activity was completely abolished, indicating full separation-of-function.

### *Prdm9*^EP^ and *Prdm9*^EK^ mutations abolish the H3K4 and H3K36 trimethylation activities of full-length PRDM9 *in vitro*

To characterize the effects of *Prdm9*^EP^ and *Prdm9*^EK^ on full-length PRDM9 *in vitro*, we introduced these mutations into a cDNA clone encoding human *PRDM9*. We then expressed and partially purified full-length wild-type PRDM9, PRDM9^EP^ and PRDM9^EK^, and repeated the HMT assays described above. In stark contrast to the isolated PR/SET domains, both mutations completely abolished the H3K4 and H3K36 methyltransferase activities of full-length PRDM9 *in vitro* (Fig. 2).

### *Prdm9*^EP^ and *Prdm9*^EK^ confer severe catalytic hypomorphism *in vivo*

We used CRISPR/Cas9 gene editing to introduce the *Prdm9*^EP^ and *Prdm9*^EK^ mutations in mice on the B6 genetic background. Some of the *in vivo* effects of these mutations were reported elsewhere (Bhattacharyya *et al*. 2019; Powers *et al*. 2020), and here we use those data for direct comparison between the two. After breeding these mutations to homozygosity, we performed ChIP-seq for H3K4me3 and H3K36me3 in spermatocytes isolated from homozygous 14-day-old males. The resulting ChIP-seq data show severe reductions in both number and intensity of PRDM9-dependent H3K4me3 peaks (Table 1, Fig. 3A-F); however, in contrast to the effects of these mutations on the full-length protein *in vitro* (Fig. 2), histone methyltransferase functions are neither separated nor abolished by either mutation. There is no effect on PRDM9-independent peaks (Fig. 3A, G). *Prdm9*^EP^ and *Prdm9*^EK^ show 4.98-fold and 5.57-fold reductions in peak number, and 2.84-fold and 3.56-fold reductions in peak intensity at detectable H3K4me3 peaks, respectively (Table 1). The effect of PRDM9^EK^ on H3K4me3 peaks is slightly more severe *in vivo* than that of PRDM9^EP^ (Table 1, Fig. 3C-F).

**Table 1:** Effect of *Prdm9*^EP^ and *Prdm9*^EK^ homozygosity on H3K4me3 and H3K36me3 activity at known PRDM9-dependent H3K4me3 peaks in spermatocytes. This table compares the number and intensity of detectable H3K4me3 and H3K36me3 ChIP-seq peaks at known PRDM9-dependent H3K4me3 peaks (n = 18,838), in wild-type B6 spermatocytes and spermatocytes homozygous for *Prdm9*^EP^ and *Prdm9*^EK^. To eliminate false positives due to the nature of H3K36me3 ChIP-seq data, detectable H3K36me3 peaks were restricted to those which also had a detectable H3K4me3 peak.

| **H3K4me3** | Detectable PRDM9-dependent Peaks | Fold reduction vs. wild-type B6 | Mean read count In detectable peaks (RPM) | Fold reduction vs. wild-type B6 |
| --- | --- | --- | --- | --- |
| B6 | 18,427 | - | 3.67 | - |
| *Prdm9*<sup>EP</sup> | 3,703 | 4.98 | 1.29 | 2.84 |
| *Prdm9*<sup>EK</sup> | 3,307 | 5.57 | 1.03 | 3.56 |
| H3K4me3 + H3K36me3 |  |  |  |  |
| B6 | 6,894 | - | 1.69 | - |
| *Prdm9*<sup>EP</sup> | 412 | 16.7 | 1.22 | 1.39 |
| *Prdm9*<sup>EK</sup> | 243 | 28.4 | 0.995 | 1.70 |

**Figure 3:**
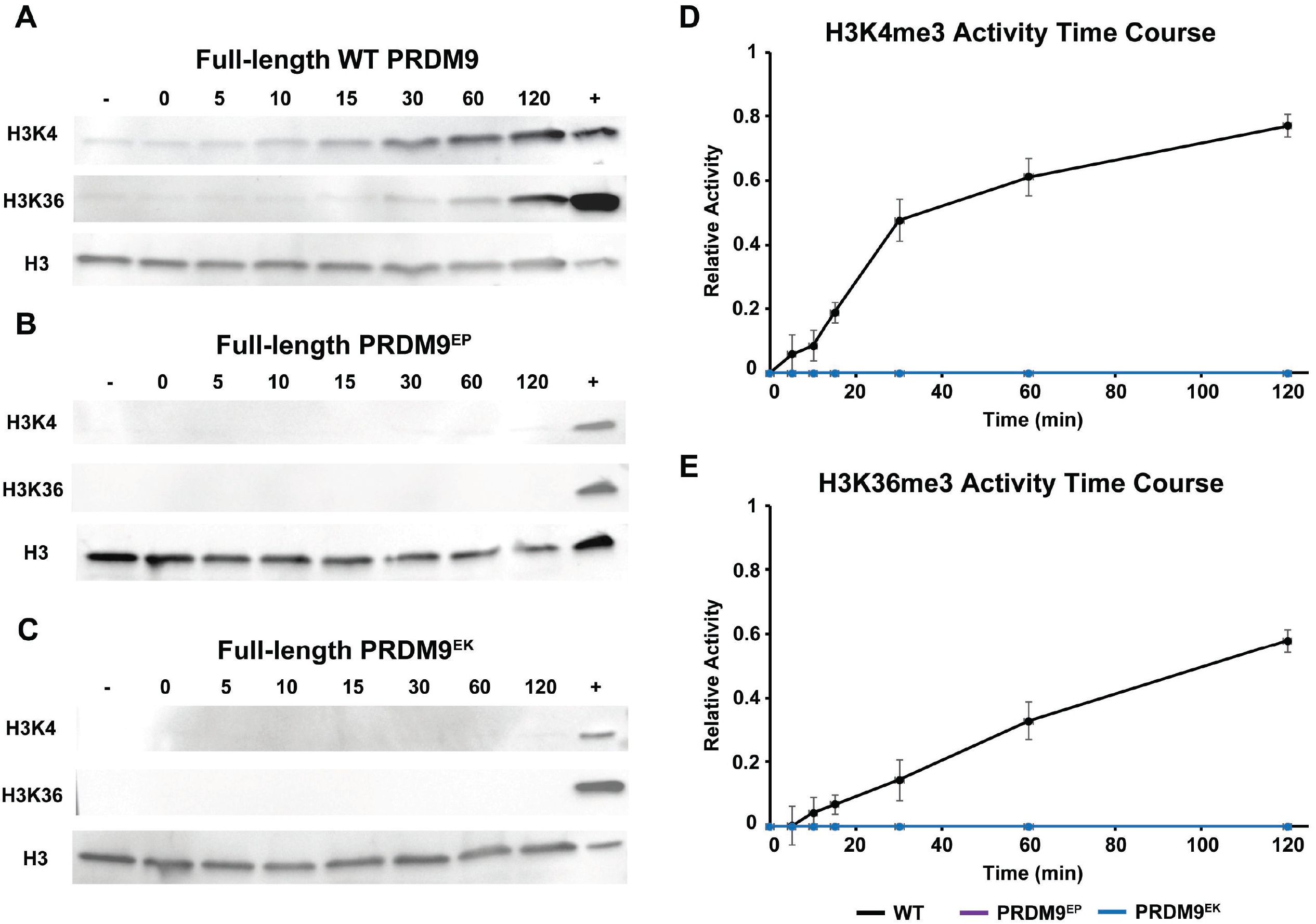
Effects of PRDM9^EP^ and PRDM9^EK^ homozygosity *in vivo*. (**A-B**) Genome browser snapshot of H3K4me3 and H3K36me3 in wild-type B6, PRDM9^EP^, and PRDM9^EK^ spermatocytes. Note the loss of both modifications at almost all hotspots, and their residual presence at one hotspot in (**B**). (**C-D**) Aggregation plots of H3K4me3 at all known PRDM9-dependent H3K4me3 peaks (n = 18,838), on a linear (**C**) and a logarithmic (**D**) scale. (**E-F**) Aggregation plots of H3K4me3 at common detectable PRDM9-dependent H3K4me3 peaks between wild-type, PRDM9^EP^, and PRDM9^EK^ (n = 2,632), on a linear (**E**) and a logarithmic (**F**) scale. (**G**) Aggregation plot of H3K4me3 at common PRDM9-independent H3K4me3 peaks between wild-type, PRDM9^EP^, and PRDM9^EK^ (n = 76,995). (**H**) Aggregation plot of H3K36me3 at all known PRDM9-dependent H3K4me3 peaks (n = 18,838). (**I**) H3K4me3 signal vs H3K36me3 signal in wild-type spermatocytes at all known PRDM9-dependent H3K4me3 peaks (n = 18,838). Sites with common detectable H3K4me3 and H3K36me3 peaks between all three genotypes (n = 118) are highlighted in black. Note the enrichment of highly active wild-type peaks in this subset.

In distinct contrast to the PRDM9-dependent H3K4me3 peaks, PRDM9-dependent H3K36me3 peaks are barely detectable in both mutants (Table 1, Fig. 3H), but still observable (Fig. 3B). Indeed, the number of detectable H3K4me3/H3K36me3 double-positive peaks is reduced 16.7-fold in *Prdm9*^EP^ homozygotes and 28.4-fold in *Prdm9*^EK^ homozygotes. The intensities of the H3K36me3 peaks that do remain, however, are not as markedly reduced as those of H3K4me3 peaks—1.39-fold and 1.7-fold in *Prdm9*^EP^ and *Prdm9*^EK^, respectively (Table 1). As shown in Fig. 3I, in wild-type spermatocytes PRDM9-dependent H3K4 and H3K36 trimethylation signals are highly correlated—the most active sites tend to be high in both. These strong wild-type peaks are enriched for detectable double-positive peaks in *Prdm9*^EP^ and *Prdm9*^EK^ (Fig. 3I). This is consistent with the expected effect if *Prdm9*^EP^ and *Prdm9*^EK^ confer lower catalytic activity *in vivo*.

### *Prdm9*^EP^ and *Prdm9*^EK^ straddle the threshold for female fertility in the B6 genetic background

In males, homozygosity for either *Prdm9*^EP^ or *Prdm9*^EK^ causes infertility with complete meiotic arrest, phenocopying the *Prdm9*-null condition (Bhattacharyya *et al*. 2019; Powers *et al*. 2020). Assessment of female fertility, however, yielded an unexpected result. As we reported previously, female *Prdm9*^EP^ homozygotes are sub-fertile, producing variable numbers of litters in natural matings (data reproduced in Table 2A-B) (Powers *et al*. 2020). In contrast, none of the female *Prdm9*^EK^ homozygotes we tested produced any pups (Table 2C). Thus, the slight reduction in PRDM9 catalytic activity exhibited by *Prdm9*^EK^ relative to *Prdm9*^EP^ *in vivo* is sufficient to abolish female fertility, in contrast to that seen in *Prdm9*^EP^ homozygotes.

**Table 2:**
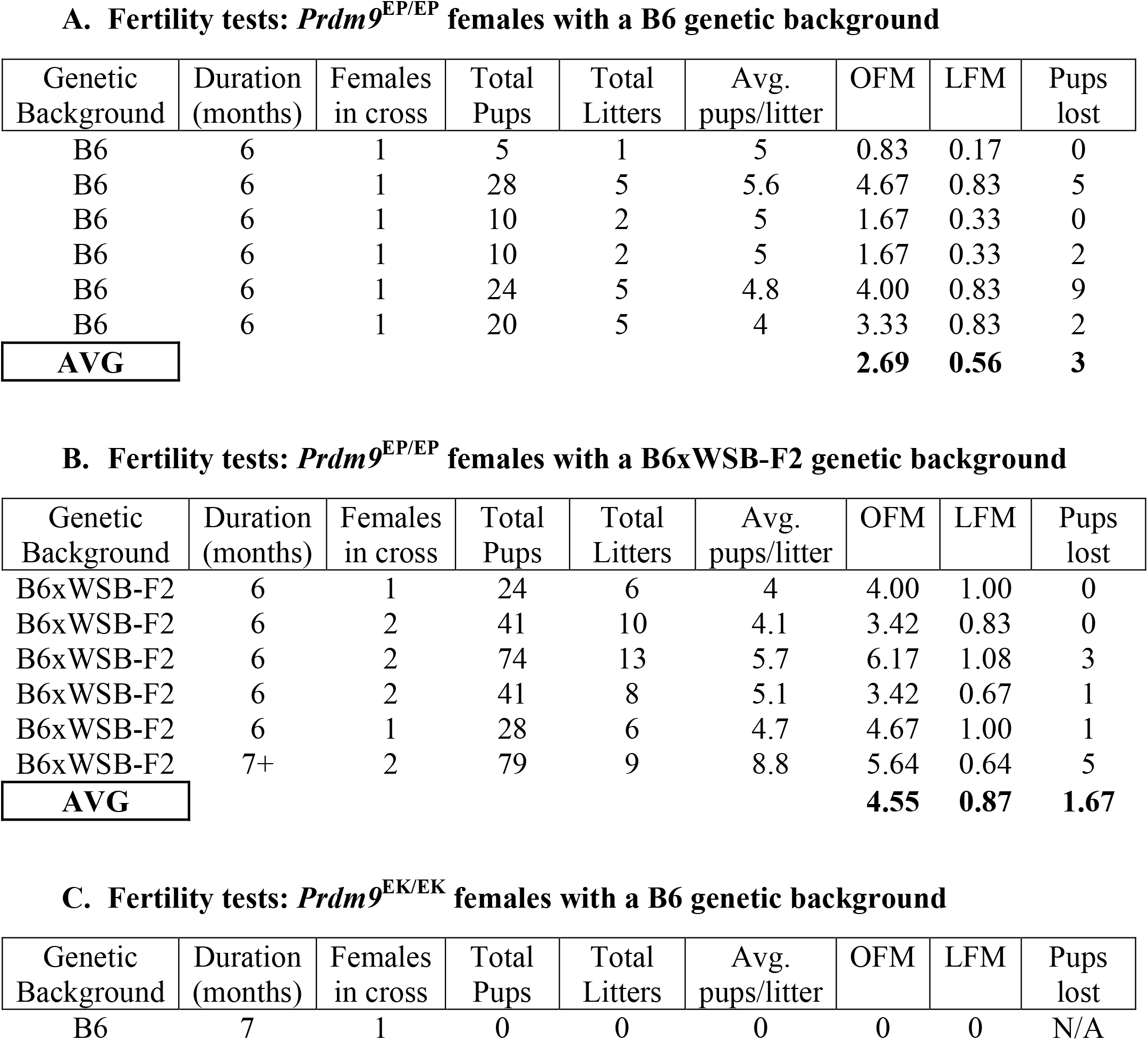
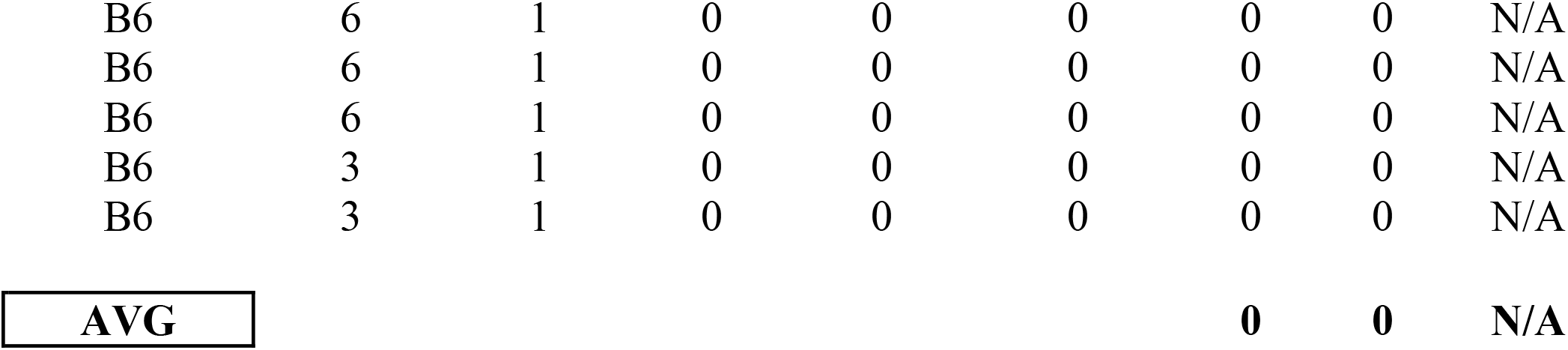
*Prdm9*^EP/EP^ and *Prdm9*^EK/EK^ female fertility tests. (**A-B**) We previously reported fertility data for *Prdm9*^EP/EP^ females elsewhere; we reproduce it here for comparison with fertility data for *Prdm9*^EK/EK^ females (**C**). Females were crossed at 5 weeks of age with a sexually mature (>8 weeks), wild-type B6 male, in pairs or trios as noted. Productive crosses were allowed to continue for at least six months, and longer if necessary until the females stopped producing offspring. Unproductive crosses were allowed to continue at least three months. The number of pups in each cross that died before wean are noted in the last column; all other offspring appeared grossly normal and healthy at wean. All offspring in the B6 background, and the offspring from at least 3 litters per cross in the B6xWSB-F2 background, were genotyped for *Prdm9*^EP^ to confirm maternal homozygosity. As expected, all genotyped offspring were heterozygous for *Prdm9*^EP^. OFM: offspring per female per month; LFM: litters per female per month.

## Discussion

*Prdm9* was identified as the mammalian recombination activator in 2010 (Baudat *et al*. 2010; Berg *et al*. 2010; Parvanov *et al*. 2010). Since then, it has amassed a degree of fame as its many curious properties have come to light. It is unique among mammalian histone modifying enzymes in several respects. First, it is the only known gene that specifically combines the KRAB, PR/SET, and ZnF functional domains. Each of these domains has unique properties. The KRAB domain binds proteins that help to open local chromatin (Parvanov *et al*. 2017; Tian *et al*. 2021). The PR/SET domain has the intrinsic capacity to *de novo* trimethylate two different lysine residues, thereby creating a unique epigenetic signature at recombination hotspots (Hayashi *et al*. 2005; Baker *et al*. 2014; Eram *et al*. 2014; Powers *et al*. 2016). The ZnF DNA-binding domain is among the most rapidly evolving protein-coding regions known in vertebrates, in contrast to other ZnF proteins where these domains are mostly conserved (Najafabadi *et al*. 2015; Wolf *et al*. 2020). Second, it is the only mammalian hybrid sterility gene identified to date (Mihola *et al*. 2009; Bhattacharyya *et al*. 2013; Lustyk *et al*. 2019). Third, its loss was thought to lead to unconditional infertility, but the growing list of exceptions to this rule suggests modifiers and different degrees of dependence on PRDM9 for successful meiosis among mammalian species, subspecies, and populations (Narasimhan *et al*. 2016; Mihola *et al*. 2019; Powers *et al*. 2020). This study now adds to understanding of the unique functions of PRDM9 in two ways: first, by pointing out the interdependence of PRDM9 domains *in vivo* and *in vitro*, and second, by elaborating the molecular and phenotypic effects of these two unique mutations, we provide new information of theoretical and potentially practical significance.

The structure of *Prdm9* suggests a modular protein: three well-separated domains with theoretically distinct functions. The catalytic PR/SET domain and the DNA-binding ZnF domain, which retain their functions *in vitro* when isolated from the rest of the protein, had indeed seemed particularly autonomous (Billings *et al*. 2013; Eram *et al*. 2014; Walker *et al*. 2015; Powers *et al*. 2016). The disparity of the effects of these mutations on the isolated PR/SET domain versus full-length PRDM9, however, shows this not to be the case. Clearly, the tertiary structure of full-length PRDM9 imposes an enzymatic context on the PR/SET domain that dramatically alters its response to catalytic mutations. In retrospect, this pseudo-modularity is perhaps not surprising in a protein encoded by such an evolutionary labile gene. *Prdm9* belongs to the KRAB-ZnF family of transcription factors; these genes are generally characterized by rapid evolution via gene duplications and repurposing for new functions through subsequent ZnF array diversification (Lupo *et al*. 2013; Imbeault *et al*. 2017). In contrast, *Prdm9* evolves a multitude of alleles of the same gene while preserving its unique function as a recombination regulator (Parvanov *et al*. 2010; Hinch *et al*. 2011; Buard *et al*. 2014; Kono *et al*. 2014).

Evolutionary analysis indicates that PRDM9-directed recombination may be present in lower taxa, and that it may have been lost in several vertebrate lineages, as it was lost in the canid lineage (Baker *et al*. 2017). A structurally complete version of *Prdm9* is present in most placental mammals and such organisms as turtles, snakes, lizards, and some fish, while partial versions missing one or more functional domains are present in many taxa. A complete version of *Prdm9* can be found in such early vertebrates as jawless fish and coelacanth, suggesting that its absence from other taxa resulted from repeated loss or repurposing of *Prdm9* (Baker *et al*. 2017). In a protein whose functions change frequently over evolutionary time, functional domains are expected to merge or separate frequently, and sometimes incompletely. Such may be the case with PRDM9; despite the modularity suggested by its structure, functional autonomy of its catalytic domain evidently has not happened.

Apart from using these two mutations as cautionary examples against assuming domain modularity when engineering enzymes, it is instructive to contemplate the results of these enzymatic manipulations. First, the *in vivo* effects of these mutations differ from those in both the isolated PR/SET domains and the full-length proteins *in vitro*. This is not entirely unexpected, because PRDM9 does not operate in a vacuum during meiosis. Unlike the situation *in vitro*, PRDM9 is bound to DNA when it trimethylates H3K4 and H3K36 *in vivo*. Additionally, multiple other proteins have been found to interact with PRDM9 during meiosis, including cohesin complexes containing the meiosis-specific protein STAG3 (Parvanov *et al*. 2017; Bhattacharyya *et al*. 2019). The RNA-binding protein EWSR1 (Tian *et al*. 2021) and the chromatin remodeling factor HELLS (Spruce *et al*. 2020) also interact with PRDM9, and both are required for efficient PRDM9-dependent catalytic activity *in vivo*. These complex interactions have not yet been placed in a temporal time-course, and, moreover, could re-orient either the PR/SET domain or the substrate, allowing for the residual activity we observe with full-length PRDM9^EP^ and PRDM9^EK^ *in vivo*, but not *in vitro*. Second, these two mutations each alter the same residue: glutamate 360 to proline and lysine, respectively. The mutations were deliberately designed based on a published crystal structure of the PR/SET domain with docked histone tail peptides (Eram *et al*. 2014). We chose proline and lysine on the principles of changing the local shape of the binding pocket and reversing the charge of the residue, respectively. Interestingly, changing the shape (Glu>Pro) was more effective than charge reversal (Glu>Lys) in separating the two methyltransferase functions. In fact, we did create a PR/SET domain that, when cloned separately, can trimethylate H3K4, but not H3K36 *de novo*. If this domain retains the same capacity when expressed alone *in vivo*, it might be possible to artificially trimethylate H3K4 at selected genomic sites by tethering it to a DNA-binding domain or to inactivated Cas9.

The unexpected female fertility phenotype of *Prdm9*^EP^ added another dimension to the hypomorphic nature of the two mutations. The phenotypic contrast between *Prdm9*^EP^ and *Prdm9*^EK^ makes these two mutations especially fascinating, as they differ only slightly in severity (Table 1, Fig. 3C-F). As we demonstrated previously, in females homozygous for PRDM9^EP^, meiotic DSBs occur at both PRDM9-dependent and PRDM9-independent sites. In some of the offspring of these females, PRDM9-independent DSBs gave rise to crossovers, indicating the presence of a PRDM9-independent crossover pathway in female mice (Powers *et al*. 2020). However, this pathway is not sufficient for successful meiosis in B6 females in the absence of *Prdm9*. The fact that *Prdm9*^EP^, but not *Prdm9*^EK^, is compatible with some fertility in B6 females suggests the possibility of a minimum threshold of PRDM9-dependent meiotic DSBs, or some other complex phenotype modification. There is evidence that inefficient repair of PRDM9-independent meiotic DSBs contributes to meiotic arrest in *Prdm9*-null animals, and this has been speculated to be its primary cause. In females, DNA damage surveillance unequivocally plays a role, as removing the checkpoint kinase *Chk2* permits marginal rescue of female fertility in *Prdm9*-null B6 mice—albeit with a high incidence of pregnancy loss and a high perinatal death rate (Powers *et al*. 2020). If *Prdm9*-null meiotic arrest is indeed due to inefficient DSB repair, our results suggest a model in which oocytes with ‘methyltransferase-comatose’ PRDM9 can complete meiosis if enough meiotic DSBs are PRDM9-dependent, allowing the oocyte to evade DNA damage checkpoint activation. We propose that in the case of *Prdm9*^EP^ homozygotes, a minority of oocytes complete meiosis, but that in the case of *Prdm9*^EK^ homozygotes, the slightly lower catalytic activity of PRDM9^EK^ shifts the balance too far in favor of inefficiently repaired PRDM9-independent DSBs, and few or no oocytes evade the checkpoint. A key difference between PRDM9-dependent and PRDM9-independent DSBs is the presence of H3K36me3 at the former, but not the latter. Since H3K36me3 is known to promote homologous recombination repair in somatic cells (Aymard *et al*. 2014; Pfister *et al*. 2014), one might speculate that inefficient repair at ectopic DSBs is due to its absence, and that H3K36 trimethylation activity could be a determinant of whether an oocyte passes the checkpoint.

*Prdm9*^EP^ and *Prdm9*^EK^ straddle the female fertility threshold for loss of PRDM9 functionality in B6 mice. Notably, as we previously observed, female fertility in *Prdm9*^EP^ homozygotes is higher in F2 crosses between B6 and WSB/EiJ (WSB), another *domesticus* strain, indicating an effect of genetic modifiers and/or hybrid vigor (Powers *et al*. 2020). Examination of the effects of these mutations in other genetic backgrounds and in outbred mice would be an interesting approach to determine the nature of threshold fertility. It is possible, for instance, that the activity of PRDM9^EP^ is not sufficient for female fertility in some strains, and that the lower activity PRDM9^EK^ is sufficient for female fertility in others. A particularly interesting case would be that of PWD/Ph, a *musculus* strain in which *Prdm9*-null *males* are able to produce small amounts of sperm, while the *Prdm9*-null females are infertile (Mihola *et al*. 2019). This strain appears not to follow the trend of female robustness to loss of PRDM9 function seen in *domesticus* and *castaneus* strains, and thus the effect of *Prdm9*^EP^ and *Prdm9*^EK^ on both spermatogenesis and oogenesis in these mice would be informative to assess. Catalytic *Prdm9* hypomorphs such as these, and *Prdm9*^EP^ in particular, may also have some practical utility for genetic mapping. Because it allows for healthy offspring with PRDM9-independent crossovers in *domesticus* strains, *Prdm9*^EP^ might permit recombination events within ‘cold spots’ refractory to PRDM9-dependent recombination.

In summary, the *Prdm9*^EP^ and *Prdm9*^EK^ mutations have revealed informative insights into the catalytic properties of PRDM9. Their dramatically different effects on the isolated PR/SET domain vs. the full-length protein show that PRDM9 does not exhibit autonomous modular function despite a structure that suggests it might. The moderately different effects of these two mutations on the full-length protein *in vitro* vs. *in vivo* suggest a regulatory and/or determinative role for proteins that interact with PRDM9 at the sites of hotspots activation. The different phenotypic effects of these two mutations reveal the minimum threshold of PRDM9-dependent catalytic activity for female fertility in the B6 genetic background. As intriguing as these discoveries are, PRDM9 likely has more peculiarities yet to be uncovered, and these mutations will continue to be useful for further study of this exceptional and fascinating gene.

## Data Availability

Strains and plasmids are available upon request. All sequence data newly reported in this study are available at the Gene Expression Omnibus (GEO) under accession number GSE181479 (reviewer access token: epqnwwqgvnaftyd).

## Acknowledgements

We would like to thank the Genome Technologies and Genetic Engineering Technology cores at The Jackson Laboratory for their invaluable expertise and assistance with this project.

